# Multi-Neuromodulator Measurements across Fronto-Striatal Network Areas of the Behaving Macaque using Solid-Phase Microextraction

**DOI:** 10.1101/534651

**Authors:** Seyed-Alireza Hassani, Sofia Lendor, Ezel Boyaci, Janusz Pawliszyn, Thilo Womelsdorf

## Abstract

Different neuromodulators rarely act independent from each other to modify neural processes but are instead co-released, gated, or modulated. To understand this interdependence of neuromodulators and their collective influence on local circuits during different brain states, it is necessary to reliably extract local concentrations of multiple neuromodulators in vivo. Here we describe results using solid phase microextraction (SPME), a method providing sensitive, multi-neuromodulator measurements. SPME is a sampling method that is coupled with mass spectrometry to quantify collected analytes. Reliable measurements of glutamate, dopamine, acetylcholine and choline were made simultaneously within frontal cortex and striatum of two macaque monkeys (*Macaca mulatta*) during goal-directed behavior. We find glutamate concentrations several orders of magnitude higher than acetylcholine and dopamine in all brain regions. Dopamine was reliably detected in the striatum at tenfold higher concentrations than acetylcholine. Acetylcholine and choline concentrations were detected with high consistency across brain areas, within monkeys and between monkeys. These findings illustrate that SPME microprobes provide a versatile novel tool to characterize multiple neuromodulators across different brain areas in vivo to understand the interdependence and co-variation of neuromodulators during goal directed behavior. Such data will be important to better distinguish between different behavioral states and characterize dysfunctional brain states that may be evident in psychiatric disorders.

**New and Noteworthy:** Our manuscript reports a reliable and sensitive novel method for measuring the absolute concentrations of glutamate, acetylcholine, choline, dopamine and serotonin in brain circuits in-vivo. We show that this method reliably samples multiple neurochemicals in three brain areas simultaneously while nonhuman primates are engaged in goal directed behavior. We further describe how the methodology we describe here may be used by electrophysiologists as a low barrier to entry tool for measuring multiple neurochemicals.

## Introduction

Extracellular concentrations of neuromodulators influence firing regimes, input-output relationships and neural interactions in local circuits and long-range brain networks (Marder 2012; Thiele and Bellgrove 2018), and are dysregulated in virtually all psychiatric disorders (Avery and Krichmar 2017; Millan et al. 2012). Accumulating evidence suggests that these fundamental roles of neuromodulators for circuit functioning are unlikely realized by single neuromodulators operating in isolation. Rather, neuromodulatory systems are heavily intertwined (Avery and Krichmar 2017; Gobert et al. 1998; Moehle et al. 2017), and operate simultaneously on individual cells and circuits (Arnsten et al. 2010; Hassan et al. 2015; Marder 2012; Santana et al. 2018). In each circuit, local mechanisms exert control over the release of neuromodulators from terminals of brainstem-originating projection neurons. This local control proceeds through activation of pre-synaptic glutamatergic receptors (Ghersi et al. 2003; Grilli et al. 2009; Luccini et al. 2007; Pittaluga 2016; Pittaluga et al. 2006; Wang et al. 1992). These insights suggest that an understanding of the contribution of neuromodulators to circuit functioning requires measuring, simultaneously, multiple neuromodulators in conjunction with ongoing glutamatergic neurotransmitter concentrations and action. Consistent with this conclusion single neuromodulator theories often fail to account for all observable symptoms in psychiatric diseases (Bohnen et al. 2015; Halliday et al. 2014; Remy et al. 2005).

Despite the accumulating evidence for the interdependence of neuromodulator actions, few methods exist for their simultaneous measurement in vivo and across multiple brain areas (**Table S1**; https://github.com/att-circ-contrl/SPME_paper_SI.git). Most of these existing neurochemical sensing methods allowing multi-neuromodulator sampling have a barrier to entry by requiring specialized equipment and trained experts preventing data collection by scientists who are otherwise interested in the role of endogenous and exogenous neuroactive chemicals in cognition and psychiatric disorders. Electrochemical methods such as fast scan cyclic voltammetry (FSCV) and amperometry have sub-second temporal resolution, but are limited to the measurement of a few compounds (Dale et al. 2005; Jacobs et al. 2010) and are challenging and not robust for wide-spread in vivo application in nonhuman primates yet, although several labs have recently reported success (Disney et al. 2015; König et al. 2018; Quintero et al. 2007; Schluter et al. 2014; Schwerdt et al. 2017; Vartak et al. 2017; Yoshimi et al. 2015). Imaging techniques such as positron emission tomography (PET) are also limited to the measurement of one or a few compounds simultaneously (Fisher et al. 1995). Microdialysis (MD) paired with mass spectrometry is the most commonly used method for measuring multiple neuromodulators in awake behaving animals. MD provides a data collection method which is then analyzed post hoc to identify and quantify collected analytes. It operates with a semi-permeable membrane which allows for the continuous collection of the available extracellular neuromodulators through passive diffusion, and can even be used to locally release pharmacological agents (Anderzhanova and Wotjak 2013; Buck et al. 2009; Kennedy 2013; Perry et al. 2009; Watson et al. 2006). However, MD does have several disadvantages. MD disrupts the tissue during its initial placement of the probe or a guiding cannula resulting in damage-induced release of neuromodulators that can last several hours before stable measurements become possible. Moreover, MD has low affinity for hydrophobic compounds and comparatively broad spatial and temporal resolution that is in the range of 200-400 μm in diameter and 10-20 minutes, respectively. These values are dependent on the surface area of the permeable membrane, the exact method of MD, flow rate, resolution of detection methods for analytes of interest, tissue tortuosity and more (Anderzhanova and Wotjak 2013; Kennedy 2013; Watson et al. 2006).

Here, we set out to address some of these limitations with a novel protocol for measuring multiple neuromodulators in vivo in discrete 20 minute intervals using probes optimized for solid phase microextraction (SPME) (Pawliszyn 2000, 2012). SPME probes are thin (200 μm) wires of arbitrary length coated with an inert porous polymeric matrix using biocompatible binder on one end where molecules with appropriate size and affinity migrate via passive diffusion and are retained by weak intermolecular interactions (*see* Methods). SPME provides an alternate method for data collection which can then be analyzed by tools such as mass spectrometers. This method has been shown to extract in neural tissue dynamic changes in dopamine (DA) and serotonin (5- HT) levels with comparable precision to MD (Cudjoe et al. 2013; Cudjoe and Pawliszyn 2014). Additionally, due to the similarity of SPME probes to commonly used microelectrodes in electrophysiological recordings, relatively minor adjustments will allow for the adaptation of conventional microelectrode driving systems for SPME use. This, combined with post collection analysis through standard chemistry facilities makes SPME an attractive and easy-to-use tool for electrophysiology labs.

SPME has the potential to be a powerful new tool to compliment the mentioned methods well suited for neurochemical profiling that spans both multiple neuromodulators as well as multiple brain regions simultaneously. Such data will allow for global observation of slow neuromodulator dynamics that could better inform our hypotheses and help relate global neuromodulator levels to electrophysiology and behavior.

Thus, the ability of SPME to report major neuromodulators as well as glutamate and GABA were tested in two behaving rhesus macaques. Probes were repeatedly and simultaneously inserted into two cortical regions and the striatum to observe inter-areal differences between extracellular neuromodulator concentrations. We found that extracellular concentrations of glutamate, dopamine, acetylcholine and choline could be reliably distinguished and differed systematically between brain regions.

## Methods

### Animals

Data was collected from two 8 year-old male rhesus macaques (*Macaca mulatta*) weighing 8-12 kg. All animal care and experimental protocols were approved by the York University Animal Care Committee and were in accordance with the Canadian Council on Animal Care guidelines. Details regarding the experimental setup, recording procedures, and reconstruction of recording sites have been described previously (Oemisch et al. 2015). Briefly, animals were implanted with a 20 mm by 28 mm recording chamber over the frontal region of the right hemisphere guided by stereotaxic coordinates (Paxinos et al. 2000) and MR images. The animals were seated in a custom made primate chair and head stabilized with their eyes 65cm away from a 21’ LCD monitor refreshed at 85 Hz. Eye traces were collected by a video-based eye-tracking system (Eyelink 1000 Osgoode, Ontario, Canada, 500 Hz sampling rate). The animals were engaged in an over-trained attention tasks in which they would use saccadic eye movements to acquire juice reward (Fig S1; https://github.com/att-circ-contrl/SPME_paper_Sl.git). The specifics of the task are described elsewhere (Hassani et al. 2017). Both animals showed stable performance and acquired similar reward volumes on all recording days. In both tasks, stimulus presentation and reward delivery was controlled through MonkeyLogic (http://www.brown.edu/Research/monkeylogic/).

### SPME protocol and Fabrication of SPME Probes

We provide a visual overview of the complete SPME protocol used here in **Fig S2** (https://github.com/att-circ-contrl/SPME_paper_Sl.git) and delineate the chemical materials, LC-MS/MS analysis, and detailed quantitation of neuromodulators in the ***Supplementary Materials*** (https://github.com/att-circ-contrl/SPME_paper_Sl.git) and in a companion paper (Lendor et al. 2019). All measurements used miniaturized SPME probes that were manufactured by repeated dip-coating of stainless steel wires in a suspension of extracting phase in a binder. This general procedure reported previously (Gómez-Ríos et al. 2017), has been modified to fit the purpose of *in vivo* brain sampling (Lendor et al. 2019). The 3 mm long tip of the wire was acid-etched down to approx. 100 μm to create a recession capable of accommodating thicker layer of coating without significantly increasing the total probe diameter. The extracting phase was an in-house synthesized hydrophilic-lipophilic balance polymer functionalized with benzenesulfonic acid to introduce strong cation exchange properties (HLB-SCX). The monodispersed polymeric particles with diameter of approx. 1 μm were suspended in the binder consisting of 7% polyacrylonitrile dissolved in N,N-dimethylformide (w/v), ensuring the particles-to-binder ratio at 15% (w/v). The extracting phase suspension was prepared one day before the probe coating, homogenized by sonication and stirred at 800 rpm overnight. Several layers of coating were deposited on the modified wires, resulting in total coating thickness of ≈50 μm and total probe diameter of 195 μm. The average pore size of the extracting phase was measured at 1.2nm. Before the use, the probes were cleaned with the mixture of methanol, acetonitrile and 2-propanol (50:25:25, v/v/v), activated in the mixture of methanol and water (50:50. v/v) and sterilized in steam for 15 min at 121°C. During the sampling, SPME probes were inserted into sheathing cannulas to protect the coating and prevent it from extracting compounds on the way to the target brain area. The cannulas underwent the same cleaning and sterilization procedure as the probes and then the lengths of both components of the SPME assembly were adjusted to 60-70 and 70-80 mm for the probes and cannulas, respectively.

### MRI Guided Electrophysiological Mapping of Target Tissue

The anatomical coordinates of the brain regions of interest were first identified through 3 T MR images. The MR images were then verified with extracellular electrophysiological recordings of the target areas, which provided the gray and white matter boundaries for the cortical sites and the dorsal most aspect of the head of the caudate nucleus. Tungsten microelectrodes were 200 μm thick with an impedance of 1-2 MΩ. All electrodes, SPME probes and their accompanying guiding cannulas were driven down into the brain and later out using software-controlled precision microdrives (Neuronitek, ON, Canada). Electrodes were connected to a multichannel acquisition processor (Neuralynx Digital Lynx system, Inc., Bozeman, Montana, USA) which was used for data amplification, filtering and acquisition of spiking activity. Spiking activity was obtained by applying a 600-8000 Hz bandpass filter, with further amplification and digitization at a 32 KHz sampling rate. For every recording day, electrodes were lowered until the first detection of spiking activity (indicative of gray matter) at the depth suggested by the MR images.

### SPME Sampling and Post-Processing Procedures

All three SPME assemblies (example of sampling using one assembly in **Fig 1A-C**) were simultaneously driven to 200 μm above the point of first spiking detection. SPME assemblies were located ~1 mm away from the electrode penetration location. Then, only the SPME probes were inserted 3 mm into the areas of interest. On average macaque cortical thickness is only 2 mm while the head of the caudate at the point of sampling was well over 3 mm. An extraction duration of 20 min was selected to reflect stable extraction time profiles for all compounds of interest. The kinetics of target analyte extraction, for example glutamate and dopamine, can be visualized in vitro through extraction time profiles (Lendor et al. 2019) (**Fig 1D-E**). Brain homogenate was spiked with known concentrations of target analytes and four replicates were collected during which multiple probes were placed in the homogenate and sequentially removed at reported times after initial insertion and quantified (LC-MS/MS procedure described below). These extraction time profiles reflect the quantifiable concentration of collected analyte over time, which is captured in sequence by a linear “kinetic” regime, a “dynamic” regime and an “equilibrium” regime over extraction time. These labels are used to describe the various rates of analyte adsorption by the SPME fibre, where the kinetic regime reflects the fastest adsorbtion of the target analyte, the equilibrium regime reflects a period of only marginal further analyte collection and an intermediate dynamic regime (Ouyang and Pawliszyn 2007). These extraction kinetics provide information about the time it takes until equilibrium is reached in a concentration independent manner that is helpful in estimating a lower limit for extraction times to collect quantifiable analyte concentrations. After the 20 min extraction event, all SPME probes were driven back 3 mm into the guiding cannulas and all SPME assemblies were withdrawn from the brain (**Fig 1A-C**). The microdrives were then removed from the chamber to enable unclamping of the SPME probes, a brief wash and then storage in glass vials surrounded by dry ice until placing them into a −80°C freezer. One entire sampling event (one extraction in 3 different brain areas together with assembling the SPME probes and cannulas, driving into and out of brain, washing and preparing for storage) was performed within 50 min, except for the first sampling event in each sampling day. The first sampling event, with the area identification using electrophysiology recording, was performed within 75 min. The removal of the SPME microprobe from the gray matter and the positioning of a new SPME microprobe in the same location limits the temporal continuity of sampling. We believe that further optimizing of the SPME microprobe switching procedure with e.g. pre-loaded SPME probes will allow replacing SPME fibres within 2-10 min. Alternatively, SPME microprobes could be used in spatially separate but adjacent guiding tubes (separation of ~300μm), which would allow to switch sampling from one to other probe without temporal delays.

**Fig 1.**
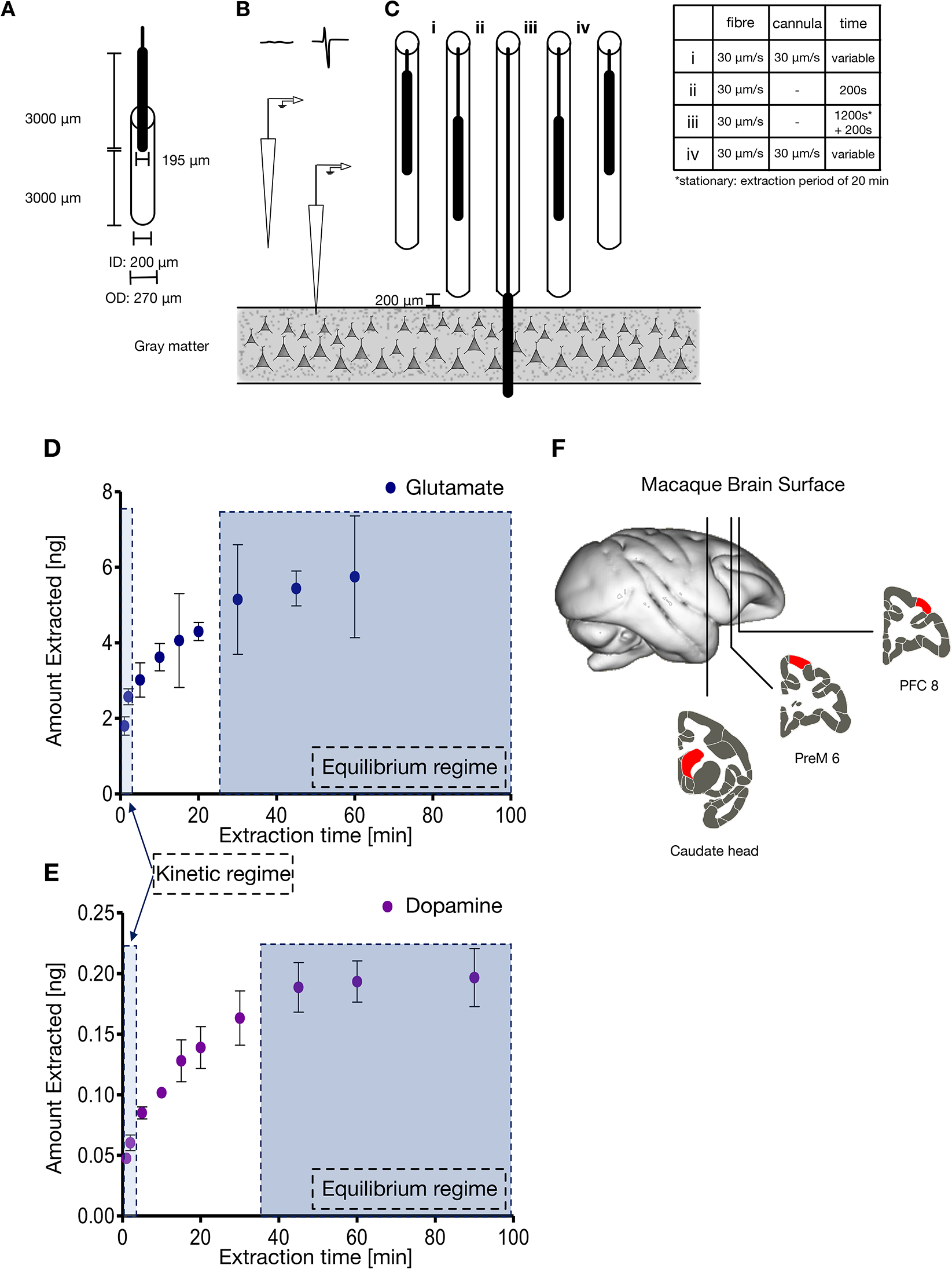
SPME sampling procedure, extraction time profiles and measurement locations. A) Dimensions of the SPME probe and the accompanying cannula. There was a 3000 μm buffer from the opening of the cannula and the start of the SPME coating. The SPME coating spanned 195 μm at the tip. The SPME coating made up the terminal 3000 μm of the entire probe and was placed in a cannula with an internal diameter of 200 μm and an external diameter of 270 μm. (B) Tungsten microelectrodes were lowered into the brain guided by 3 T magnetic resonance (MR) images in order to map the depth at which detectable spiking was observed matching expectations from the MR images. (C) SPME probes and their accompanying cannulas in all brain areas were then simultaneously lowered to 200 μm above the point of first observable firing in the target brain region and the SPME probes were lowered to expose only the 3000 μm SPME coating. After 20 minutes of extraction, the SPME probes were then retracted 3000 μm back into the cannula and the probes plus their accompanying cannulas were removed from the brain. The table inset into the figure provides the velocities and times of each probe and cannula as they transitioned between the extraction steps. (D) In vitro extraction quantities of Glutamate in ng and (E) Dopamine in ng over time utilized to select in vivo extraction times. The linear “kinetic” regime and “equilibrium” regime are labelled with the dynamic range being in between. (F) Anatomical locations of SPME sampling events in the right hemisphere of two Rhesus Macaques. Target brain regions are highlighted in red and include two cortical regions, prefrontal cortex (PFC) area 8 and premotor cortex (PreM) area 6 as well as a subcortical region: the head of the caudate.

### Neuromodulator detection and quantitation

Detection and quantitation of neuromodulators followed previously established procedures described in the **Supplementary Materials** (https://github.com/att-circ-contrl/SPME_paper_Sl.git). In brief, SPME probes were defrosted and desorbed in an aqueous solution containing water, acetonitrile, methanol, formic acid and deuterated isotopologues of target neuromodulators as internal standards (IS). Chromatographic separation of target compounds was conducted as previously reported (Cudjoe and Pawliszyn 2014). Tandem mass spectrometry (MS/MS) was performed with electrospray ionization with two MS/MS transitions for each neuromodulator and one for each IS (Lendor et al. 2019).

Stock solutions of all target neuromodulators were prepared and used to generate standard calibration curves and to calculate the amounts of neuromodulator extracted by each experimental SPME probe. Conversion of the amount of neuromodulator extracted to in-brain concentrations was done using matrix-matched external calibrations. LOD is defined as 3 times the signal obtained from blank samples.

## Results

During all measurements animals were engaged in a cognitively demanding task with stable behavioral performance to minimize state dependent fluctuations of basal extracellular concentrations of neuromodulators. Three brain regions were selected to provide a sample of extracellular neuromodulatory concentrations. Two cortical regions: prefrontal cortex (PFC) area 8, premotor cortex (PreMC) area 6 and the head of the caudate nucleus (CD) were selected (**Fig 1F**). In 3 daily sessions, we sampled 3 times per session simultaneously from all three areas. Monkey As had an additional 3 days of recording with one sample collected on each day. Overall we collected and analyzed 12 probes in monkey As and 9 probes in monkey Ke. We were specifically interested in major neuromodulators and neurotransmitters but successfully measured other compounds such as amino acids (e.g. glutamine, taurine, phenylalanine etc) that we do not discuss here.

To allow comparison of the SPME extraction results to those typically reported in microdialysis studies, we calculated the relative change in measured concentrations across the three successive sampling events per session pooled across monkeys to enhance the statistical power of the analysis (**Fig 2**). Relative to the end of the first sampling event, the second and third sampling events started after 40 minutes and 100 minutes respectively. We expected that the variability of measurements (indexed as standard error of the median) is comparable to repeatedly measured microdialysis of an identical, active brain state. We found that measured concentrations did not change significantly across sampling events for any compound area combination (Wilcoxon rank sum test; **Fig 2**).

**Fig 2.**
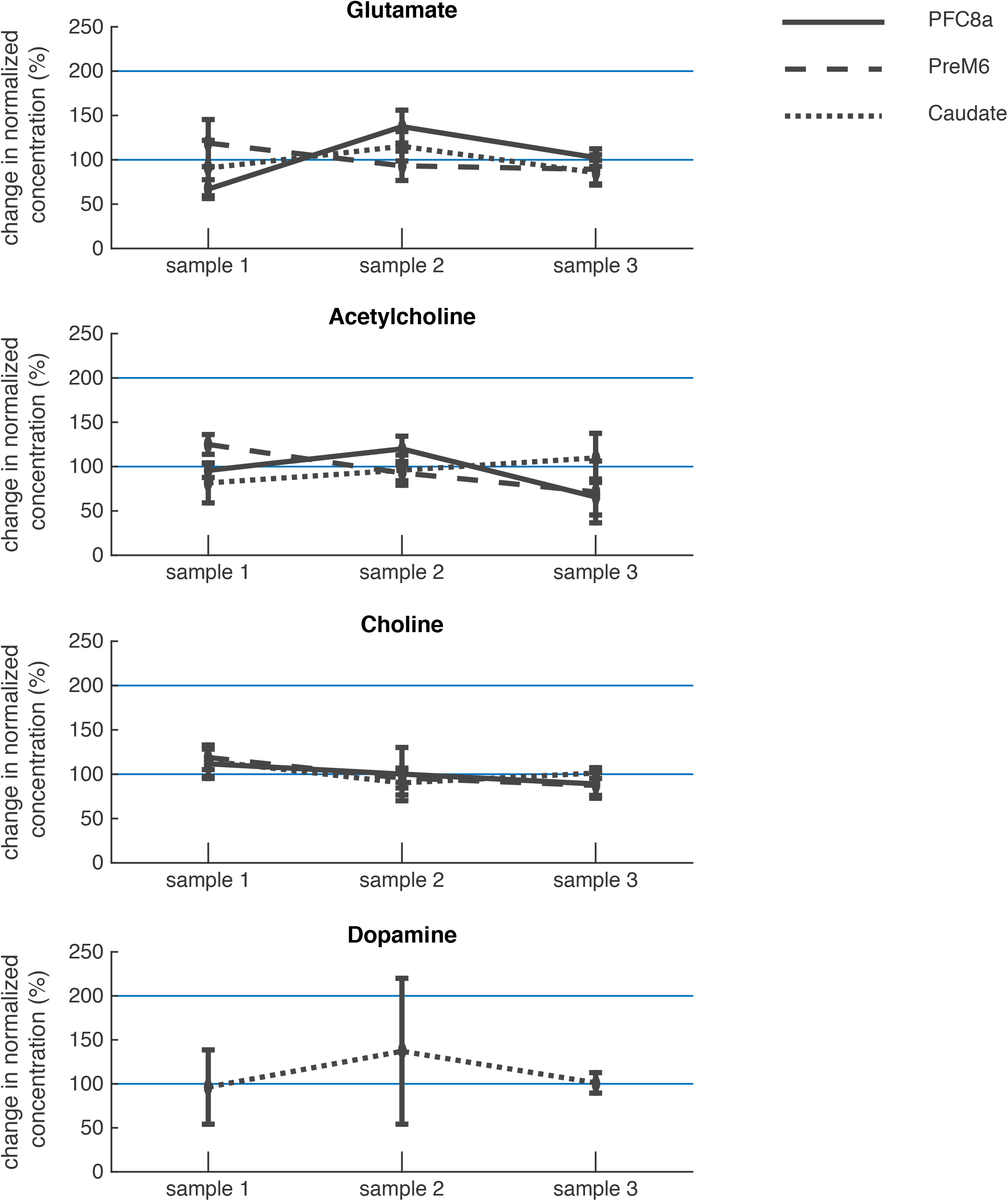
Changes in SPME sampling over time. Changes in the normalized concentrations of the observed neuromodulators across daily sampling sessions in each sampled brain region. The median concentration of (A) Glutamate (B) acetylcholine (C) choline and dopamine (D) with standard error of the median plotted for each sampling time and location. Data was normalization for each subject separately and for the respective neurochemical and area combinations. Sampling locations are represented by different lines: solid line for the prefrontal cortex, dashed line for the premotor cortex and dotted line for the caudate nucleus. Sampling events were 20 minutes in length and the time between the end of one sampling event to the start of the next sampling event was 40 minutes (STD 2 minutes) making the second sampling event 40 minutes and the third 100 minutes from the initial measurement. No significant change was observed for any neuromodulator in any brain region (Wilcoxon rank sum test). Note that a single prefrontal choline data point was excluded in this analysis for being >4 standard deviations from the median (data point is present in **Fig 3a**).

We found that four target neurotransmitters and neuromodulators were reliably detected in each animal: glutamate, dopamine, acetylcholine, and choline. Serotonin was also detected on several probes but always near the limit of detection (LOD) and therefore was excluded from analyses here (Lendor et al. 2019). Glutamate concentrations were several orders of magnitude higher than all other observed compounds of interest in all areas and both animals. Relative to glutamate, choline concentrations were >15 times lower, DA concentrations were >700 lower and ACh concentrations were >8300 times lower (**Fig 3**).

**Figure 3:**
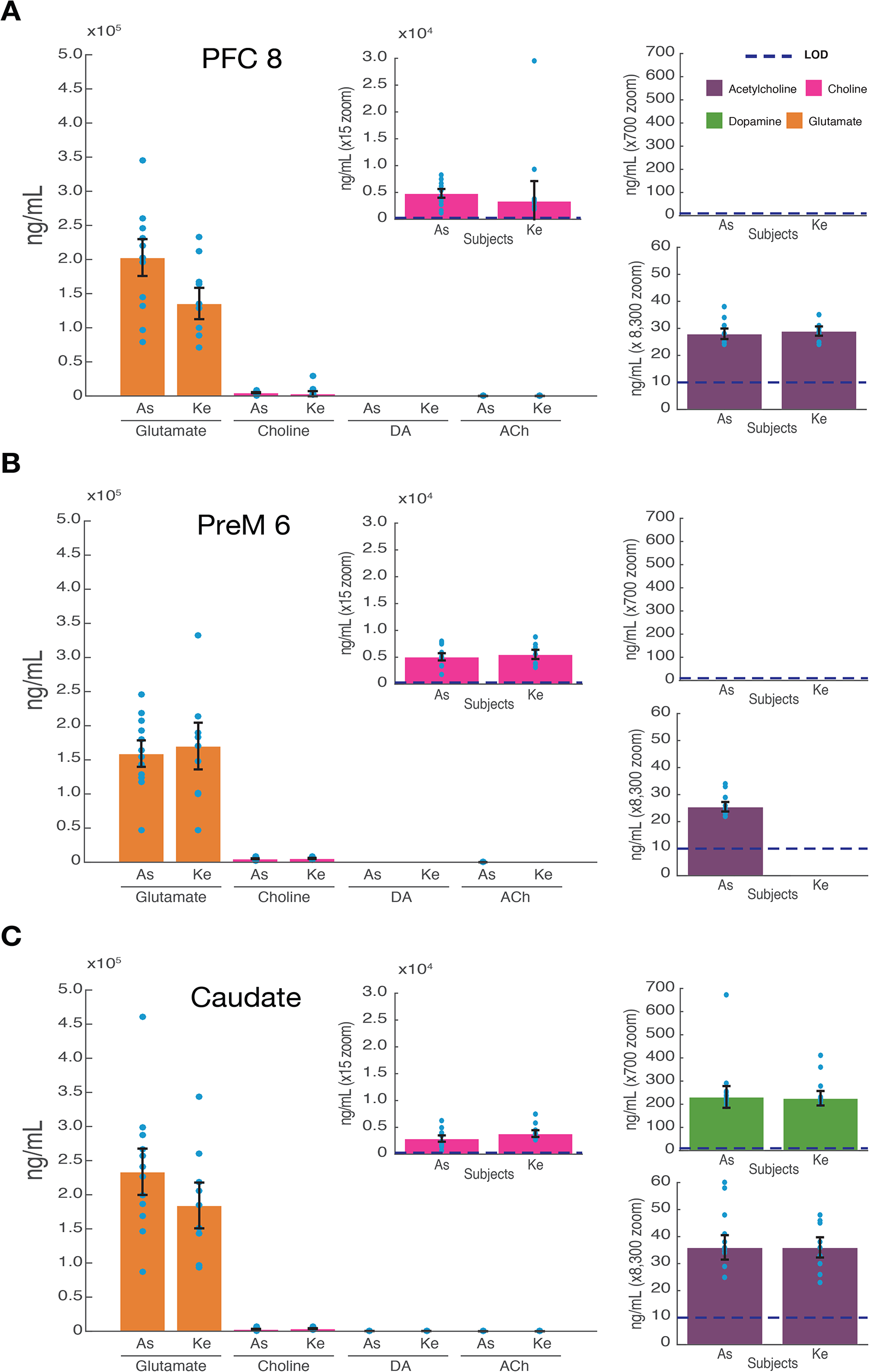
SPME sampling of the macaque brain. SPME probes extract from the cortex (PFC and PreM6) as well as the caudate of two adult macaques after 20 minutes of extraction within the respective brain region. Median and standard error of the median of n= 12 probes for animal As and n= 9 probes for animal Ke with no data excluded. Four neuromodulators were reliably measured: glutamate, dopamine, acetylcholine, and choline (orange, green, purple, red respectively). All brain regions had high concentrations of glutamate and relatively much lower concentrations of acetylcholine and choline extracted (with the exception of animal Ke PreM6 lacking detectable acetylcholine). The only tested region to yield measurable dopamine was the caudate. The dashed blue line depicts the limit of detection (LOD).

Glutamate concentrations (across areas), measured as median ± standard error, ranged from 159,147 (±19,395) to 233,659 ng/mL (±33,917) in monkey As, and from 135,523 (±22,945) to 184,333 ng/mL (±33,516) in monkey Ke. In both monkeys, glutamate concentrations were highest in the head of the caudate, which was significantly different from glutamate concentrations in the premotor cortex in monkey As (kruskal-wallis test with Tukey’s HSD correction, p = 0.048) but not in monkey Ke (kruskal-wallis test with Tukey’s HSD correction, p = 0.847, n.s.). Glutamate concentrations between the caudate and PFC, as well as the PFC and premotor cortex were not significantly different for either animal. Comparisons between monkeys showed no significant differences between the glutamate concentrations in any of the measured areas (**Fig 3**).

In addition to glutamate, dopamine could be reliably detected in both monkeys in the caudate at concentrations of 232 ng/mL (±47) in monkey As, and 226 ng/mL (±31) in monkey Ke. These concentrations were similar between monkeys (Wilcoxon rank sum test p =1, n.s.). Dopamine was not found above detection limits in frontal cortex.

Acetylcholine measurements in the three brain areas ranged from (across areas) 25 (±2) to 36 ng/mL (±5) in monkey As, and from 29 (±2) to 36 ng/mL (±4) in monkey Ke. In monkey As, extracellular ACh concentrations between the caudate and premotor cortex were significantly different (kruskal-wallis test with Tukey’s HSD correction, p = 0.008). Cortical areas were not significantly different from one another (kruskal-wallis test with Tukey’s HSD correction, p = 0.476, n.s.) nor was there a significant difference between the caudate and PFC (kruskal-wallis test with Tukey’s HSD correction, p = 0.145, n.s.). In monkey Ke, only one cortical region had concentrations of ACh above detection limits, which approached significant difference to the concentrations measured in the caudate (Wilcoxon rank sum test, p = 0.08). Measurements of extracted ACh in the pre-motor cortex of monkey Ke were consistently below detection limits (**Fig 3**).

We also measured choline which is a main product of the enzymatic breakdown of the highly regulated ACh and which is a main indicator of attentional modulation of cholinergic activity in frontal cortex (Parikh et al. 2007). Choline concentrations ranged across areas from 2,914 (±590) to 5,078 ng/mL (±675) in monkey As, and 3,408 (±3,705) to 5,533 ng/mL (±860) in monkey Ke. Choline concentrations between the caudate and premotor cortex were significantly different in monkey As (kruskal-wallis test with Tukey’s HSD correction, p = 0.030), but not in monkey Ke (kruskal-wallis test with Tukey’s HSD correction, p = 0.729, n.s.). All other area comparisons were not significant for both monkeys. Comparisons of choline concentrations between monkeys were not significant in any of the measured areas.

Overall, serotonin concentrations were not observed as reliably as the other reported neurochemicals and therefore were excluded from the main analyses. However, in monkey As, serotonin was observed near the LOD in the prefrontal cortex as well as the caudate in a subset of probes. Within the 9 probes placed in both areas, serotonin was detected in 22% of samples (2/9 probes) in the prefrontal cortex and 33% of samples (3/9 probes) in the caudate nucleus. Serotonin observations ranged from 149 ng/mL to 232 ng/mL with a median of 171 ng/mL (±18.5 ng/mL; standard error of the median) with a LOD of 100 ng/mL. The proximity of the measurements to the LOD suggest that the other probes likely collected concentrations of serotonin below the detection threshold. No serotonin observations were made in monkey Ke.

## Discussion

We demonstrated the reliable measurements of glutamate, dopamine, acetylcholine and choline simultaneously within cortical and subcortical regions of awake and behaving macaques using coated microprobes optimized for solid phase microextraction. Glutamate concentrations were several orders of magnitude higher than dopamine, acetylcholine and choline across brain regions. Extracellular concentrations of choline, dopamine and acetylcholine were detected at >15, >700, and >8,300 times lower than glutamate respectively. Dopamine was readily detected in the caudate but not observed at detectable concentrations in the cortical regions measured. Acetylcholine concentrations showed a statistical trend of being different between the caudate and measured cortical areas and with high consistency between monkeys. Choline concentrations, a product of acetylcholine degradation as well as a precursor for its synthesis, were negatively correlated with acetylcholine (R = −0.355; p = 0.01). Together, these findings provide new and rare insights about the neurochemical circuit profiles during an active brain state in three areas of the primate fronto-striatal network.

The high consistency of measured concentrations within animals, between brain areas and between monkeys suggests that SPME microprobes could provide a versatile neuro-technique for understanding how variations of the neurochemical milieu relate to local and long-range circuit operations and ultimately cognitive functioning.

### Extracellular Concentrations of Glutamate, Dopamine, Acetylcholine and Choline

Our results revealed particularly high levels of glutamate within all measured brain regions (PFC, PreMC, and CD) relative to other neuromodulators. This finding is consistent with its ubiquitous role as major excitatory neurotransmitter. This holds true even in the caudate which consists exclusively of GABAergic neurons, but receives glutamatergic inputs from the cortex (Haber and Knutson 2010). Tonic concentrations of glutamate within the striatum have been previously reported to be comparable to that of the hippocampus, prefrontal cortex and other cortical areas due to glutamatergic inputs, extra-synaptic release and glial release (Ascenzo et al. 2006; Moghaddam 1993; Moussawi et al. 2011). In contrast, classical neuromodulators such as DA and ACh, are released in extracellular space through volume transmission where they are highly regulated and may bind to receptors on multiple neurons (Santello et al. 2012; Schwartz and Javitch 2013; Syková 2004; Vizi 2000). This regulation may help explain the relatively low observed concentrations of DA and ACh and may be a large source of extracellular choline through the enzymatic breakdown of ACh.

Extracellular DA was measured well above the detection threshold in the striatum. In combination with glutamatergic and cholinergic concentrations, our results show promise in the application of SPME to further our understanding of the relationship between glutamatergic inputs to the striatum and the neuromodulatory ACh and DA signals impinging on striatal circuits. Such circuits are strongly involved in hypo-and hyperkinetic diseases (Halliday et al. 2014; Millan et al. 2012). Dopaminergic signals did not exceed the detection threshold in either cortical regions.

The presence of DA in the PFC is well documented (Goldman-Rakic et al. 1990; Vijayraghavan et al. 2007; Watanabe et al. 1997) and we thus conclude that DA most likely has failed to exceed our detection limit. The difference between the detection success of DA in the caudate vs PFC and PreM cortex may be due to differences in available DA, release concentrations and dynamics (Holloway et al. 2019). A possible explanation for the lack of detectable cortical DA is the susceptibility of catecholamines to oxidation. This may be partially addressed by faster coupling of sample collection to quantitation by MS.

Few previous studies have documented ACh concentrations in vivo in non-human primates due to its rapid breakdown by acetylcholinesterase and the challenges of neurochemical testing in primates (Kreitzer 2009; Schwartz and Javitch 2013; Zapata et al. 2013). ACh plays a major role in organizing local circuits, deployment of attention, locomotion and reward through different receptors present in cortical and subcortical regions (Granger et al. 2016; Howe et al. 2017; Moehle et al. 2017). It has been difficult to discern which ACh concentration corresponds to efficient endogenous circuit operations. Our finding of measurements in all sampling locations except for premotor cortex in monkey Ke, is therefore a promising starting point for future studies comparing ACh concentrations, in conjunction with other neuromodulators, during different cognitive states. The fast turn-around of ACh due to AChE however, likely leads to an underestimation of ACh concentrations by SPME. Although AChE cannot traverse the porous SPME coating due to its size (~7.6nm diameter), preventing any enzymatic degradation of trapped ACh, the dynamic equilibration process continuously leads to a transfer of ACh out of the SPME coating following the concentration gradient of the rapidly depleted ACh into the extracellular space as discussed in the case of calibration in the coating approach (Alam and Pawliszyn 2016; Zhang et al. 2007a).

Amongst other sources, choline is created as a byproduct of ACh degradation by AChE. Choline is also a precursor for the synthesis of ACh. Given this relationship, a negative correlation between choline concentrations and ACh is expected which we indeed observed in our dataset. Despite the difference in the measured concentrations of choline and acetylcholine, other studies with similar differences have demonstrated the relationship of ACh and choline to behavior within the cortex (Parikh et al. 2007). Future studies may focus on quantifying this relationship across a wider range of brain states.

Many factors may contribute to variation in measured concentrations of neuromodulators including behavioral state, the specific brain region measured and individual differences amongst others. Importantly, different methods may yield different estimates of absolute extracellular concentrations. For example, comparable measurements of extracellular glutamate via MD or voltammetry as opposed to electrophysiological estimates can differ by orders of magnitude and may be attributed to differences in probe size, or sensitivity to different sources of glutamate (Moussawi et al. 2011).

### Reliable Measurement of Individual Differences of State Specific Neuromodulatory Tone

Our results suggest that SPME probes provide reliable and sensitive measurements in consecutive sampling events within an experimental session and between sessions on consecutive days. These results were obtained while we controlled brain states across measurement events by engaging animals in a cognitive task. This experimental control could have contributed to the comparable extracellular levels of neuromodulators from sample to sample within *and* between days from similar brain locations (**Fig 2**) (Marder et al. 2014; McGinley et al. 2015; Reimer et al. 2014, 2016). The reproducibility of multiple measurements at the same anatomical location within and between days suggests that the SPME penetrations did not significantly disturb tissue (Cudjoe et al. 2013). This conclusion contrasts with reported experience from microdialysis measurements, where the initial placement of the probe or guiding cannula causes transient measurement instabilities that can require several hours of settling time before reliable, steady state neuromodulator concentrations are measurable (Kodama et al. 2002, 2015, 2017; Moussawi et al. 2011; Watanabe et al. 1997). These waiting periods must be taken into account when designing microdialysis experiments for in vivo tracking of neuromodulators during behavioral tasks in awake and restrained animals. Furthermore, the reliability of observations for the reported neurochemicals with SPME is extremely high and outperforms many reported microdialysis studies. We present 100% reliability in reporting glutamate and choline concentrations in all brain regions and subjects. Acetylcholine concentrations were reported with 100% reliability in one animal and 92% (11/12 probes) in the other within the prefrontal cortex, 0% reliability (below detection threshold) in one animal and 83% (10/12 probes) in the other within the premotor cortex and 100% reliability in one animal and 92% (11/12 probes) in the other within the caudate. Dopamine concentrations were reported with 100% reliability in both animals in the caudate and were not observed above detection thresholds in the cortex.

### Qualities and Advantages of SPME

Our results illustrate several inherent advantages of using SPME to measure neurochemical profiles in brain circuits (Lendor et al. 2019). As illustrated in **Table S1** (https://github.com/att-circ-contrl/SPME_paper_SI.git), SPME complements or outperforms alternative techniques in various domains. Practically, the arbitrary length of the wire where the SPME coating is placed allows for robust placement within the brain. The narrow diameter of the SPME probe that is within the range common to electrodes for electrophysiological recordings, allows for repeated, simultaneous sampling at multiple sites without observable changes to detected extracellular neuromodulator levels using 20-minute sampling times.

#### Spatial Resolution

In principle, the spatial resolution of the SPME measurements are capable of reflecting laminar concentration gradients across areas as small as several tens of micrometers. In our protocol we coated 3mm in length with 50 μm of thickness and desorbed the entire area into an organic solvent mixture for subsequent LC-MS/MS analysis. However, with the use of other techniques, such as desorption electrospray ionization (DESI), which represents a direct-to-MS approach (Deng et al. 2014), much smaller areas of the SPME matrix may be analyzed at a time allowing for laminar or near-laminar resolution. Direct coupling of SPME fibres to MS is the preferred mode, as it provides the most rapid analysis thus preventing diffusion of compounds within the coating and smearing the imprinted layer information. If diffusion of molecules within the fibre is high, it can be prevented by, for example, the inclusion of a divider between two halves of the coating. This would allow the simultaneous measurement of separated compartments of a cortical column. The absolute spatial resolution of SPME is difficult to estimate as SPME operates through diffusion and dynamic equilibration similar to MD. As a consequence, the sampled volume will depend on sampling time, the compound’s diffusion coefficient, as well as the target tissue and its properties affecting the rate of diffusion such as tortuosity (Syková 2004).

#### Temporal Resolution

Several factors determine the appropriate extraction time using SPME. Depending on the goals of an experimenter, resolutions below 5 minutes may be achieved as the detection limit is reached within the first few minutes for most compounds tested (**Fig 1D-E**). As sampling continues, the SPME coating will eventually reach an equilibrium point with the external environment (**Fig 1D-E**). In principle, sampling times beyond a stable equilibrium point will result in the same extracted values of compounds. However, in the brain there is no single stable equilibrium point and thus, even with long sampling times, there will be some variation in extracted concentrations. Within the dynamic regime, we can still calculate environmental concentrations from which sampling occurred given knowledge of the extraction time and the dynamic range. The lower limit of this dynamic range that can be informative is determined by the coating’s physicochemical characteristics and sample’s properties and in practice also volume of desorption and sensitivity of the MS. The analytical technique’s sensitivity also practically determines the thickness of the SPME coating as thinner coatings could in principle increase temporal resolution but would extract less compounds overall requiring higher sensitivity to detect. With higher sensitivity, less time is required for the thin coating to extract sufficient amounts to exceed limits of detection and quantitation. This then means that the compound with regards to which the sensitivity is the lowest, limits the temporal resolution because it will have the slowest-to-reach detectable concentration. But as long as the detection threshold is exceeded, the extraction time profiles allow for the calculation of the equilibrium concentration even with pre-equilibrium sampling. However, making measurements in the dynamic range requires consistency of extraction times as differences in time will lead to variations of extraction yields in a range that can be estimated from the SPME extraction time profiles (**Fig 1D-E**). Given the dynamic nature of brain networks and the lack of a stable equilibrium point, we expect consistency in brain states to be one of the biggest factors in reducing replicated variability. Moreover, SPME measurements are inherently an average of the temporal dynamics of the target tissue milieu. This means that due to the bi-directional exchange of neuromodulators from the extracellular space and the SPME coating, longer periods of stable concentration gradients will be more strongly reflected in the extracted measurements.

#### Utility and Ease of Use

SPME based microprobe extraction allows for multiple simultaneous measurements that can be reliably repeated within the same measurement sessions at the same locations with no evidence of damage induced disruption of the neurochemical environment. Another critical advantage of SPME over comparable MS-analysis based methods such as MD is its ease of use and accessibility. Due to their similar size to recording microelectrodes used for electrophysiology, little adjustment is required for conducting SPME measurements using existing acute microelectrode positioning systems. Measurements in deeper structures should use an accompanying guide tube (**Fig 1C**) as travelling through non-target tissue will result in unwanted chemical collection. Beyond the collection, a chemistry core could apply LC-MS/MS to the probes allowing for relatively easy quantitation of extracellular compounds of interest.

#### Detection Methods

SPME is ultimately a sampling method, much like MD, that provides data for analytical tools such as chromatography and mass spectrometry. In fact, once SPME analytes have been desorbed from the probe into a solvent, data provided by SPME and MD are treated very much the same. Thus, the advancement in detection limits and reliability of post-hoc analytical methods utilized by MD also benefit SPME, and vice versa. However, SPME sample analysis is more reliably and conveniently coupled to LC-MS/MS as it lacks issues such as high salt content of collected dialysate that MD suffers from (Guihen and Connor 2009).

#### Extension of SPME measurements beyond classical neuromodulators

A unique advantage of SPME over alternative methods of in vivo detection of compounds within the brain is its potential affinity for hydrophobic compounds. Although MD is capable of collecting metabolomics data similarly to SPME (Anderzhanova and Wotjak 2013; Li et al. 2012; Zhang et al. 2007b), the artificial cerebra-spinal fluid that is commonly traversing the semi-permeable membrane better facilitates the collection of hydrophilic molecules. This means that molecules that cross the semi-permeable membrane and are therefore collected and quantified through LC-MS are much more likely to be hydrophilic. SPME, in contrast, provides balanced analyte coverage, which means that in principle certain SPME matrix coatings have similar affinity for both hydrophilic or hydrophobic compounds (Reyes-garcés et al. 2018). Data collected here includes other neurochemicals of interest such as amino acids (e.g. glutamine, taurine, phenylalanine, etc) that we reliably detected but do not discuss here. In practice, the detection of very polar molecules such as monoamines and catecholamines requires coating chemistry that facilitates hydrophilic compound extraction. Comparatively many lipids play important roles in intracellular signaling and have been suggested to provide biomarkers for psychiatric disorders (Tamiji and Crawford 2011; Yehuda et al. 1999), therefore making SPME a potentially versatile and comprehensive tool for brain neurochemistry studies.

### Future direction and improvements to the SPME neurochemical sensing

The next immediate steps for the continued testing of SPME’s utility as a neurochemical sampling method is to evaluate its ability to report behavioral state dependent changes in extracellular neuromodulator concentrations. Similar testing as described here will be conducted with varying, stable behavioral states such as passive engagement, active task engagement, and drowsy/sleepiness. We predict that the various brain regions tested will display different changes in neuromodulator concentrations as a function of behavioral state.

Several improvements to the protocol can be made in order to make SPME more informative. Using a more sensitive MS would reduce detection thresholds and increase sensitivity in detecting compounds not successfully measured here such as GABA, serotonin or norepinephrine. This would be very impactful, especially in the case of serotonin for reasons discussed above. Additionally, efforts could be made to improve the extracting phase synthesis and functionalization protocols and increase the extracting capabilities of SPME probes. Moreover, in order to decrease the MS background and interferences in the range of small molecules, derivatization strategy could be considered.

For GABA, the most promising strategy is post-desorption derivatization by reagent increasing the compound’s hydrophobicity resulting in better MS signal such as benzoyl chloride (Wong 2016). Catecholamine and acetylcholine detection may be improved by optimizing the post-sampling analysis pipeline. Improvements can be made at several steps in order to better preserve and quantify catecholamines: (1) faster coupling of sampling to desorption (2) trying various antioxidant solutions and other preservatives for maintaining catecholamine integrity and (3) faster analysis of the desorbed analytes. Faster coupling of sampling to desorption could be achieved through a more stream-lined process of fibre placement and retraction involving a static cannula maintained through several sampling events. This mechanism would also allow for a faster replacement of SPME probes resulting in a shorter delay between consecutive measurements.

A preliminary experiment using both spiked artificial cerebra-spinal fluid (aCSF) and lamb brain homogenate with very high, known concentrations of all target compounds yielded detectable concentrations of all target compounds indicating that the extracting phase of the SPME probe is capable of capturing all compounds of interest. Furthermore, simultaneously collected replicates displayed very little loss of collected compounds over several days within a −80°C freezer (data not shown). Various strategies were compared for storage and placement of extraction fibres in desorption solvent prior to −80°C storage seemed most reliable for many compounds and will be the storage method utilized in the future. Such findings neither support nor antagonize the suggested strategies for improving SPME yield and sensitivity in primates as detection properties within the aCSF and lamb brain homogenate may not accurately reflect those observed in vivo in primates. This is likely contributed to by the dynamic and highly regulated nature of target compounds within the extracellular space of the brain and the dynamic equilibrium between the extracting phase of the SPME probes and the extracellular environment. Further tests of the possible differences between these mediums and improvement strategies to SPME probes are subject for future studies.

### Implications for Understanding and Treating Psychiatric Disease States

Most psychiatric disorders are accompanied by neuromodulatory dysregulation (Millan et al. 2012). However, this fact is seldom studied in a multi-modulatory manner with a single or few nuclei or neurochemicals being observed at a time. This is an increasingly evident problem because models attributing symptoms of a disorder to a single neuromodulator often fall short in explaining many symptoms (Bohnen et al. 2015; Remy et al. 2005). For example, in Parkinson’s disease, outside of the well characterized dopaminergic deficits, there is evidence for deficits in noradrenergic, cholinergic and serotonergic systems as well (Halliday et al. 1990, 2014; Moehle et al. 2017). Many cognitive deficits observed within Parkinson’s disease are linked to such non-dopaminergic deficits. Such findings emphasize the need to simultaneously observe multiple neuromodulatory systems. Thus the simultaneous measurement of multiple neuromodulators, in multiple brain regions within healthy and clinical populations will allow for a better understanding of the underlying causes of symptoms and progression of psychiatric disorders (Anderzhanova and Wotjak 2015; Avery and Krichmar 2017; Millan et al. 2012).

A better understanding of psychiatric disorders through multi-modulator methods may also lead to more accurate understanding of the action of pharmacological agents. Previously, many studies aimed at identifying the locus of action of a pharmacological agent have used local injection methods such as iontophoresis or microinjections (Lapiz and Morilak 2006; Wang et al. 2007). Although highly informative about the role of neuromodulators in modulating the activity of individual cells and circuits, this approach does not allow physiologically realistic exploration of a systemically administered pharmacological agent. Pharmacological agents are often administered in some systemic fashion and even with highly specific receptor affinities, may interact with multiple neuromodulatory systems through the actions of heteroreceptors (Avery and Krichmar 2017; Gobert et al. 1998; Millan et al. 2015). Multi-modulator measurements, as described here, will allow for a better understanding of pharmacological agents, as well as provide novel insights into the development of more effective drugs or combinations of drugs to better treat the clinical population.

### Conclusion

We described a novel SPME protocol capable of simultaneous, multi-modulator measurements of multiple brain regions. Our results suggest that SPME both supplements current methods of neuromodulator detection and allows for novel measurements previously not possible for the investigation and dissection of neuromodulatory systems, their role in physiological brain processes and their modulation by pharmacological agents.

## Acknowledgments

This research was supported by grants from the Canadian Institutes of Health Research (CIHR), the Natural Sciences and Engineering Research Council of Canada (NSERC) and the Ontario

Ministry of Economic Development and Innovation (MEDI). We thank Dr. Hongying Wang for invaluable help with animal care. We thank Thermo Scientifc for providing the triple quadrupole mass spectrometer TSQ Quantiva used in this work as a loan to our laboratory. The authors thankfully acknowledge Pfizer Canada Inc., Merck Canada Inc., Quebec Consortium for Drug Discovery (CQDM), Brain Canada, and Ontario Brain Institute for the grant “Solid phase microextraction-based integrated platform for untargeted and targeted in vivo brain studies”.

## Supplementary Methods

### Chemicals, Reagents and Materials

The LC-MS-grade solvents methanol (MeOH), acetonitrile (ACN), isopropanol (IPA) and water, as well as the acetylcholinesterase inhibitor phenylmethylsulfonyl fluoride (PMSF) were purchased from Fisher Scientifc. Formic acid, polyacrylonitrile (PAN), N,N-dimethylformamide (DMF) and the standards of neurotransmitters: *γ*-aminobutyric acid (GABA), glutamic acid (Glu), acetylcholine (ACh), histamine (Hist), serotonin (5-HT), dopamine (DA) and choline (Cho) as well as their deuterated analogues were purchased from Millipore Sigma (Oakville, ON, Canada). Epinephrine (Epi), norepinephrine (NE) and their deuterated analogues were obtained from Cerilliant Corporation (Round Rock, TX, USA). Choline-D9, was purchased from Cambridge Isotope Laboratories (Tewksbury, MA, USA). The reagents used for synthesis of hydrophilic-lipophilic balance polymer particles functionalized with strong cation exchange groups, as well as compounds for preparation of PBS were purchased from Millipore-Sigma. The stainless steel wire (stainless steel grade AISI 304, 150 μm diameter) used for manufacturing of SPME probes was purchased from Unimed S.A. (Lausanne, Switzerland). The stainless steel tubing used as guiding cannulas (270 μm O.D.; 200±5 μm I.D.) was obtained from Vita Needle (Needham, MA, USA).

### LC-MS/MS Analysis

On the day of analysis, the SPME probes were defrosted and desorbed into 40 μL of water/ACN/MeOH 40:30:30 (v/v/v) mixture containing 1 *%* of formic acid and a mixture of deuterated isotopologues of targeted neuromodulators (used as internal standards, IS) at 20 ng/mL. The desorption was carried out for 1 h with agitation at 1500 rpm. The extracts were injected into the LC-MS system for targeted neurotransmitter analysis within a few hours after desorption. All experiments were carried out using an Ultimate 3000RS HPLC system coupled to TSQ Quantiva triple quadrupole mass spectrometer (Thermo Fisher Scientific, San Jose, California, USA). Data processing was performed using Thermo software Xcalibur 4.0 and Trace Finder 4.1. For chromatographic separation of the target compounds, a modified method of what is previously reported (Cudjoe and Pawliszyn 2014) was used, adjusted to include more targeted neuromodulators and their corresponding IS. A Kinetex^®^ PFP LC column (100 × 2.1 mm, 1.7 μm; Phenomenex, Torrance, CA, USA) was held at 30°C with the mobile phase flow rate at 400 μL/min. The mobile phase A consisted of water/MeOH/ACN 90:5:5 (v/v/v) with 0.1 % formic acid, and mobile phase B was ACN/water 90:10 (v/v) with 0.1% of formic acid. The chromatographic gradient was applied starting from 100% B for 1 min and increasing the aqueous mobile phase A to 100% for 3 min with convex gradient function, held for 0.5 min and subsequent linear return to initial conditions and re-equilibration for 1 min, yielding total time of 6.5 min. The injection volume was 10 μL. MS/MS analysis was performed with electrospray ionization in positive mode under selected reaction monitoring conditions, with two MS/MS transitions for each neuromodulator (quantifier and qualifier) and one for each IS.

### Quantitation of neuromodulators

Individual stock standard solutions of all targeted neuromodulators were prepared in methanol or water with 0.1% formic acid at a concentration of 1 mg/mL and stored at −80 °C for maximum of one month. In order to calculate the amounts of neuromodulators extracted by each probe, calibrator standards prepared in the same desorption solvent as real samples were analyzed in the same batch. This instrumental calibration curve was prepared in the range of 0.1-200 ng/mL by a serial dilution of the stock standard mixture of all compounds at 1 μg/mL. The IS concentration was kept constant at 20 ng/mL in all calibrators, identically as in the real samples. The amounts extracted were calculated based on linear regression equation obtained from the analytical signal (the ratio of chromatographic peak areas of analytes and their corresponding IS) plotted against the concentration.

In order to calculate the concentrations of neuromodulators in brain, matrix-matched external calibration approach was used. The surrogate matrix consisted of 2% agar gel mixed with brain homogenate in the ratio 1:1 (*v/w*). The homogenized brain tissue was earlier incubated with 1 mM PMSF for 1 h at 37°C to prevent enzymatic digestion of acetylcholine in the calibrator samples. Due to several target compounds being present in brain homogenate at high concentrations (e.g. for glutamate and choline the “blank” brain homogenate matrix doesn’t exist), their quantitation was based on signals of their deuterated isotopologues. The calibrator samples were prepared in the surrogate matrix with concentrations of neuromodulators ranging from 5 to 3000 μg/mL for the isotopically labelled compounds or from 10 to 2000 ng/mL for the remaining compounds.

The extractions were carried out with SPME probes manufactured and pre-treated identically to the probes used for *in vivo* sampling and using the same 20 min extraction time and desorption conditions as for the real samples. The amounts of neuromodulators extracted from the calibrator samples were determined in the same way as described above and plotted against concentrations of calibrators. The resulting weighted linear regression equations were applied to the amounts of neuromodulators extracted from the *in vivo* samples, yielding values of concentrations of the compounds of interest in brain.

The limits of detection (LOD) were estimated as the levels corresponding to the signal to noise ratio of 3 and were calculated based on the signal of blank calibrator sample (considered as the noise).

## Supplementary Figures

**Fig S1.**
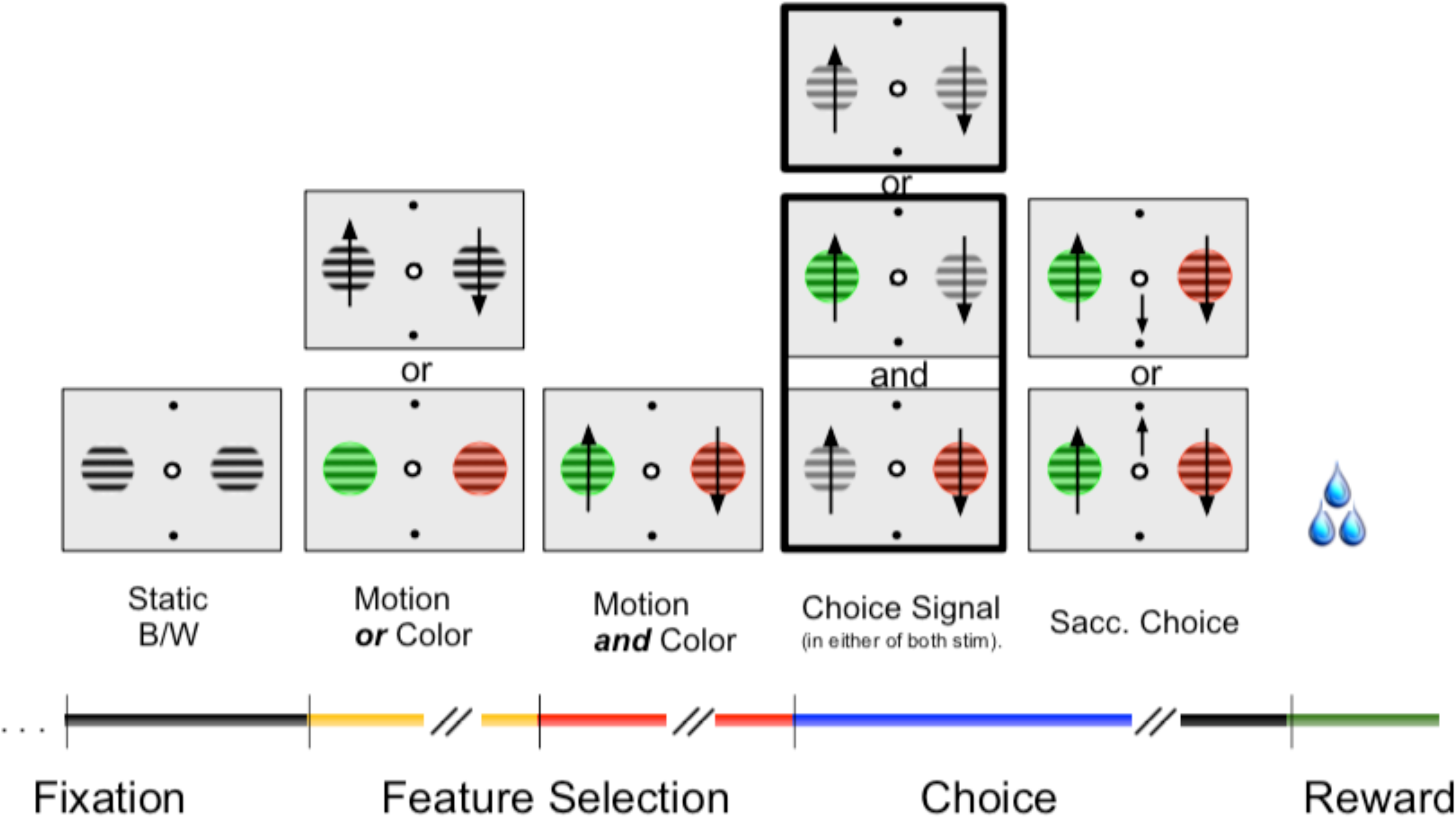
Behavioral task that the monkeys were engaged in. Briefly, the monkey was expected to fixate a central fixation point until criterion when two graded stimuli appeared. The graded stimuli acquired color and motion, of the graded stripes, features in either order. The two stimuli then either simultaneously dimmed (go-signal), or dimmed one at a time in either order. The monkey, through trial and error, identified the rewarded stimulus via its color feature which was the sole identifying feature informative of reward. The monkey was then expected to wait until the dimming of the selected stimulus and respond in the same direction as the motion of the graded stripes on the chosen stimulus. If the monkey correctly accomplished this, it would receive deterministic reward in the form of liquid juice. Monkey As was engaged in a variation of this task with reduced complexity in order to match monkey Ke in performance and reward acquisition over the sampling period. Figure reproduced from Hassani et al., 2017.

**Fig S2.**
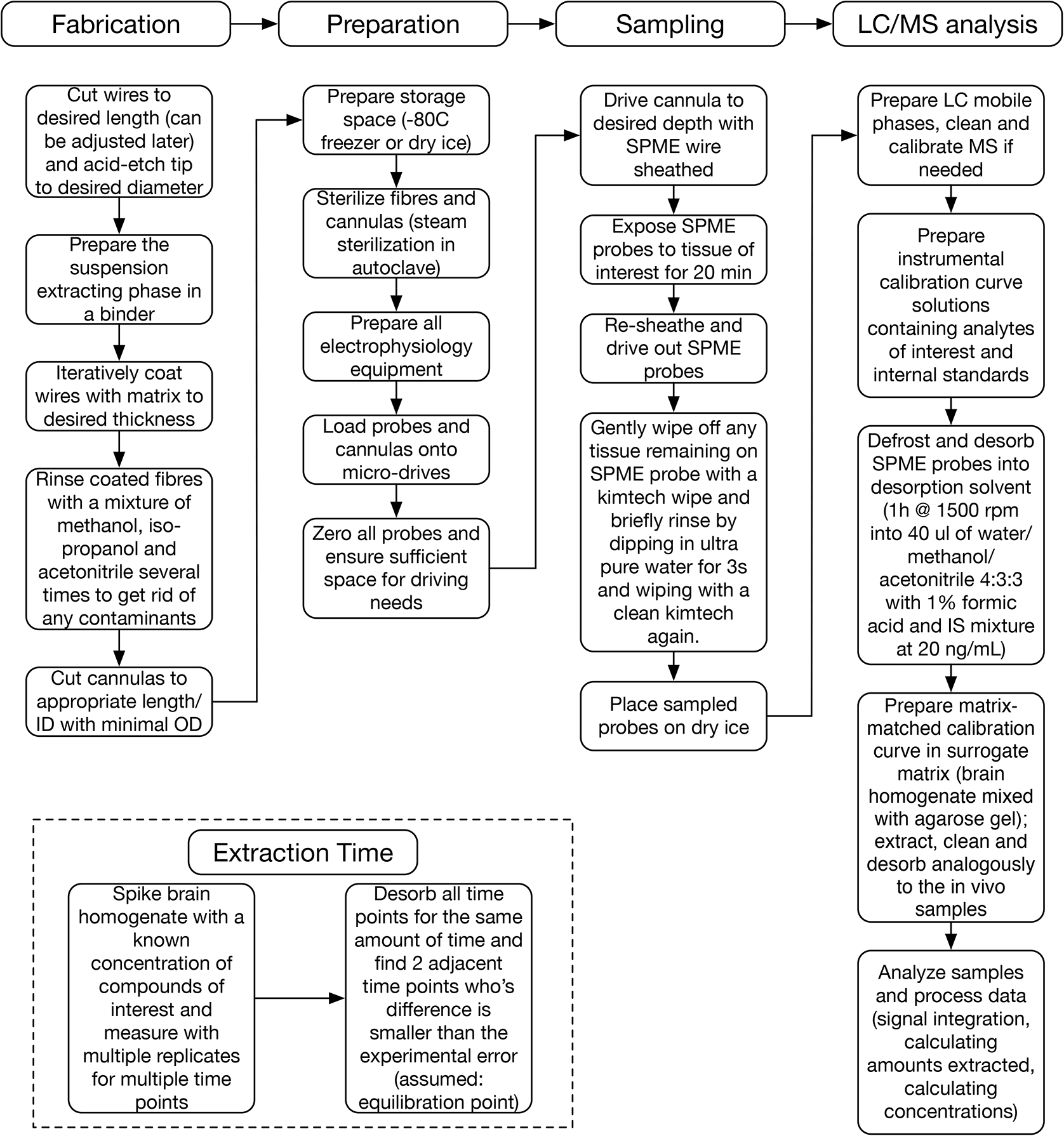
Experimental procedure from SPME probe fabrication to quantitation. Fabrication described the in-house procedure to prepare SPME probes. Preparation describes experimental setup. Sampling describes the actual data collection process. LC/MS analysis describes the chemical quantitation of collected data samples.

## Supplementary Tables

**Table S1:**
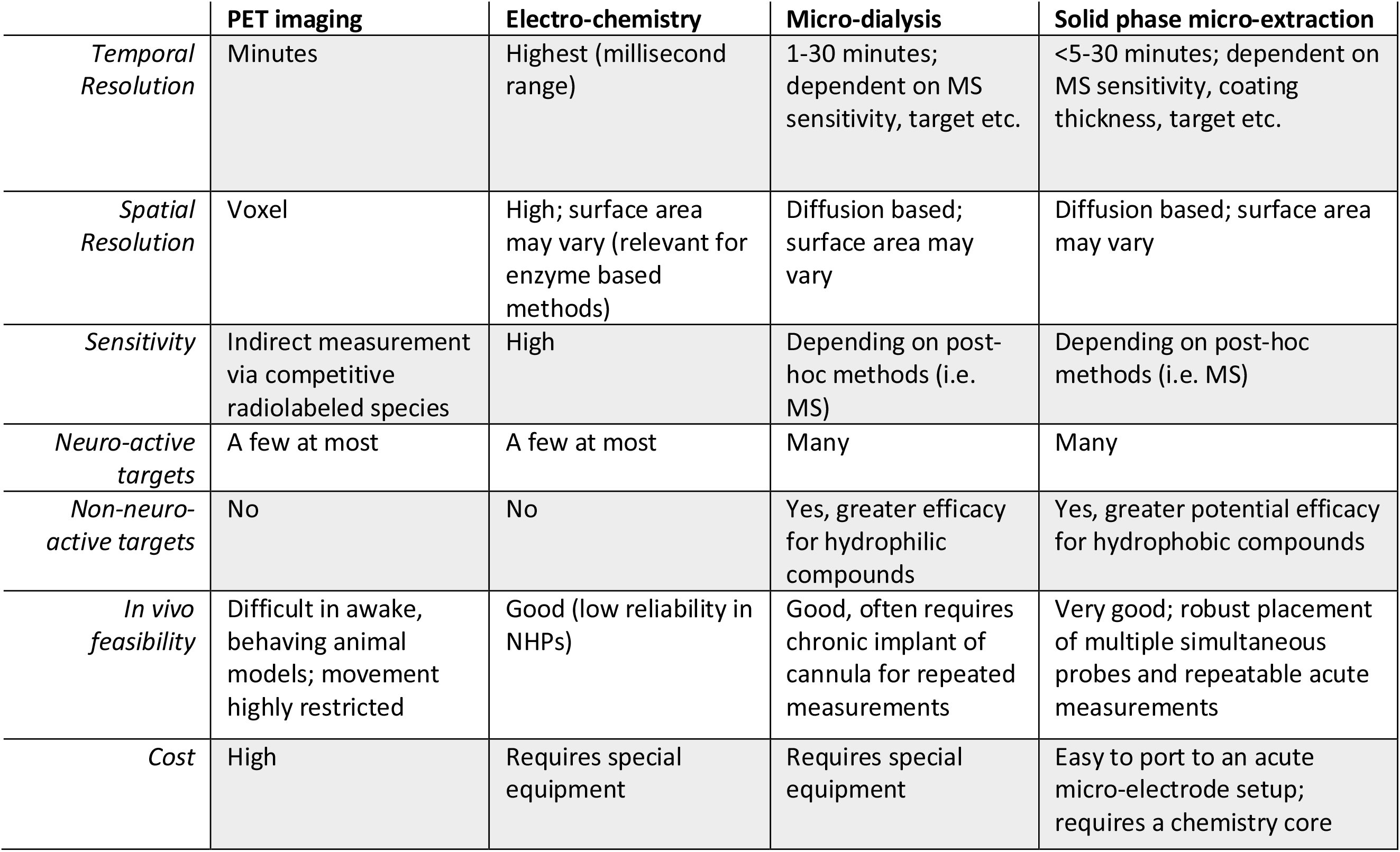
Overview of in vivo neurochemical measurement methods

**Table S1**. A comparison of methods capable of measuring single or multiple neurochemicals in vivo. Temporal resolution, spatial resolution, sensitivity, capability to measure neuro-active and non-neuro-active compounds, in vivo feasibility and cost. PET: positron emission tomography; NHP: non-human primate; MS: mass spectrometer.

